# Insights from two decades of the Student Conference on Conservation Science

**DOI:** 10.1101/819623

**Authors:** Jonas Geldmann, Helena Alves-Pinto, Tatsuya Amano, Harriet Bartlett, Alec P. Christie, Lydia Collas, Sophia C. Cooke, Roberto Correa, Imogen Cripps, Anya Doherty, Tom Finch, Emma E. Garnett, Fangyuan Hua, Julia Patricia Gordon Jones, Tim Kasoar, Douglas MacFarlane, Philip A Martin, Nibu Mukherjee, Hannah S. Mumby, Charlotte Payne, Silviu O. Petrovan, Ricardo Rocha, Kirsten Russell, Benno I. Simmons, Hannah Wauchope, Thomas A. Worthington, Rosie Trevelyan, Rhys Green, Andrew Balmford

## Abstract

Conservation science is a crisis-oriented discipline focused on delivering robust answers to reducing human impacts on nature. To explore how the field might have changed during the past two decades, we analyzed 3,245 applications for oral presentations submitted to the Student Conference on Conservation Science (SCCS) in Cambridge, UK. SCCS has been running every year since 2000, aims for global representation by providing bursaries to early-career conservationists from lower-income countries, and has never had a thematic focus, beyond conservation in the broadest sense. We found that the majority of submissions to SCCS were based on primary biological data collection from local scale field studies in the tropics, contrary to established literature which highlights gaps in tropical research. Our results showed a small increase over time in submissions framed around how nature benefits people as well as a small increase in submissions integrating social science. Our findings also suggest that students and early-career conservationists could provide pathways to increased availability of data from the tropics and for addressing well-known biases in the published literature towards wealthier countries. We hope this research will motivate efforts to support student projects, ensuring data and results are published and made publicly available.

## 1. Introduction

Conservation science focuses on understanding and reducing the negative impacts of human activities on nature, and has, from its inception, been framed as a “mission-oriented discipline” (Soulé 1985). It has its origins in biology and, as a result, its initial emphasis was on describing and explaining the distribution of biodiversity as well as the ecological and evolutionary processes shaping the diversity of life under human pressure. However, over the last few decades it has become increasingly clear that better understanding biological processes alone is insufficient in identifying robust solutions to reduce pressures on nature and the environment (Balmford and Cowling 2006; Bennett et al. 2017; Kareiva and Marvier 2012; Meine et al. 2006). This has led to the integration of the social sciences, economics, and psychology to understand the role of people in addressing conservation problems (Mace 2014; Martin et al. 2012b; Teel et al. 2018) and an interest in the motivations for conserving nature (Greenwald et al. 2013; Kareiva 2014; Kareiva and Marvier 2012; Noss et al. 2013; Soule 2013).

Even though primary data are the foundation for conservation science and management (Tewksbury et al. 2014; Wilson 2017), the proportion of published studies based on primary data collection has decreased over the past two decades, though they still represent 70% of ecological studies (Ríos-Saldaña et al. 2018). In addition, the conservation literature continues to exhibit considerable geographical biases toward wealthier, often English-speaking countries (Amano and Sutherland 2013; Martin et al. 2012a) and certain taxonomic groups (Clark and May 2002) and away from the tropics (Collen et al. 2008; Mammides et al. 2016; Meijaard et al. 2015),. These biases limit our ability to assess what conservation actions work and where.

However, analysis of trends in peer-reviewed articles can give an unrepresentative picture of the work being done on the ground (Godet and Devictor 2018). Understanding the extent to which the peer-reviewed literature is missing specific types of studies or research from certain parts of the world can help to highlight publications gaps and improve the uptake of data and experiences outside the published literature sphere. One possible pathway to address the evidence gap and entrenched biases is to analyze conference submissions. While not without possible biases of their own, conference submissions may be less vulnerable to some of the issues in the peer-reviewed literature (e.g. positive-results publication bias, English language skills) and could therefore offer an additional view of what is happening on the ground, thus helping to identify the disconnect between on-the-ground research and the published literature with the aim to utilizing the full potential of the conservation research community.

In this study, we assessed the scope, data and methods of studies submitted for presentation at the Student Conference on Conservation Science (SCCS) in Cambridge, UK using a database of >3000 applications compiled for this study. To our knowledge, SCCS is the oldest dedicated student conservation conference. Over the 20 years it has been running, it has welcomed applications from bachelor, masters and PhD students. It has never had a thematic focus but instead encourages submissions from across the diverse disciplines of conservation science. It has consistently received applicants from around the world, in part thanks to its provision of bursaries to those from lower income countries. We classified these applications to explore patterns and trends over time in what conservation students study, focusing on potential changes in framing, the types of studies conducted, the methods used, and the integration of data and ideas from the social sciences. We were particularly interested in understanding if the transition from conservation as a predominantly biological science to a more multi-disciplinary one had changed the framing around the value of nature to people or the integration of the social sciences.

## 2. Material and methods

We included 3,487 submissions for oral presentations at SCCS covering 15 individual years spanning 2002-2019. This represented the years for which we had all conference submissions, regardless of whether the applicant had subsequently been invited to present at the conference. Ethics approval was obtained through the Human Biology Research Ethics Committee, School of Biological Sciences, University of Cambridge (ref no.: PRE.2018.068). All submissions were anonymized, and e-mails were sent to all applicants asking them to reply if they did not want to be included in the study. This led to the removal of seven submissions.

The data extraction protocol and guidelines were developed prior to reviewing the submissions (Table S1). The protocol was pilot tested on a subset of submissions (n = 20) by a sub-group of reviewers and subsequently revised based on these experiences. Two workshops were conducted prior to the data extraction to explain and discuss the final protocol. In total, 25 reviewers participated in the data extraction. The conference-submissions were assigned randomly among all 25 reviewers, with each reviewer extracting data from approximately 140 abstracts. The year of submission was removed to avoid biasing the data extraction.

### 2.1 Data extraction

For each submission (title and abstract), the reviewers extracted information on the applicant (nationality, country of residence, career stage) as well as on 25 elements pertaining to the research carried out by the student. The abstracts for 2002, 2003, and 2006 consisted of a title and an abstract with no formatting requirements. For subsequent years the abstract was divided into four parts: 1) What conservation problem or question does your talk address?, 2) What were the main research methods you used?, 3) What are your most important results?, and 4) What is the relevance of your results to conservation?. The 25 elements covered research locations (e.g. country, region); study type (i.e. field, laboratory, modelling, remote sensing); and scale of study (e.g. local, national, multi-country) (see Table S1 for the full list and definitions). Where possible, answers were assigned to predefined categories (e.g. realm of study: terrestrial, marine, freshwater, coastal, or multiple). In addition, reviewers used ‘not sure’ where the abstract did not allow a clear interpretation or ‘not applicable’ where the particular questions were not relevant.

Where one or multiple species were studied, we recorded the broad taxon using 16 categories: algae, lichens, plants, fungi, arthropods, marine invertebrates, other invertebrates, fish, amphibians, reptiles, birds, mammals, other, multiple, not applicable, and not sure.

For each conference submission each reviewer assessed whether the study primarily addressed ‘Pressure’, ‘State’, or ‘Response’ following the PSR-framework of the Organisation for Economic Co-operation and Development (1993). For example, a study could examine the effect of protected areas (response) in reducing hunting (pressure) on numbers of lions (state). This was done based on an interpretation of the entire abstract. Where more than one category could apply, we used a hierarchical approach to assign a single category to each submission, where ‘response’ superseded ‘pressure’ which superseded ‘state’ - so the example above would be classed as a response study. The hierarchical approach was used to reflect the conceptual thinking behind the PSR-framework, that conservation is the human response to human pressures affecting the natural state of the world.

We extracted information on how far human dimensions were included in the studies through two questions. The first addressed whether the submission mentioned conservation benefiting people and/or the importance of involving people in conservation decisions. It was not necessary for the study to be primarily framed around the value of nature to people, only that the role of, or relevance to, people was articulated. The second addressed whether the primary focus of the study was the value nature provides to people.

We assessed whether submissions recorded biological data (e.g. species, habitats, genetics or any other data derived from a biological system) and/or socio-economic data (e.g. livelihood issues, economy/finances, attitudes, human behavior, or human behavior change). Additionally, we recorded if any data was collected by the students themselves as well as if there was any not collected by the student.

Finally, we recorded the methods for both biological (e.g. transects, camera-traps, remote sensing, interviews) and socio-economic data collection (e.g. interviews, questionnaires). For biological methods the original 20 categories (Table S1) were reduced to six: 1) field data, 2) genetic data, 3) internet/literature search, 4) audio and camera recordings, 5) remote sensing, and 6) other.

Following data extraction, 359 (11.1%) submissions were selected for kappa analysis to test the inter-reviewer variability in data extraction. This was done by randomly selecting 10% of the conference-submissions of each reviewer to be re-reviewed by a different randomly selected reviewer. For the years 2002 and 2003 we assessed 20% of each year following the same procedure. Kappa analysis was conducted on all questions individually and on overall agreement. Based on this, questions with a Cohen’s kappa score below 0.6 (weak agreement) were not included in the analysis (McHugh 2012). The average Cohen’s kappa for all included questions was 0.78 (S.E. = 0.07, min = 0.64, max = 0.87, Table S2). Only the identification of main threat (Cohen’s kappa = 0.21) did not meet this criterion. The years 2002 and 2003 were assessed separately leading to the exclusion of the Pressure-State-Response questions for those years (Cohen’s kappa = 0.40).

### 2.2 Analysis

Prior to calculations of proportions, all empty fields, ‘not applicable’, and ‘not sure’ were removed. Thus, the number of responses for each year varies across analyses. For questions where we assessed proportional changes over time, we used beta-regression, suitable for proportions, to model the proportion as the dependent variable and year as a continuous independent variable. All analyses were carried out in R 3.5.1 (R Development Core Team 2019).

## 3. Results

### 3.1 Geographical and taxonomic focus

We assessed 3,245 submissions after removing 235 that had been submitted but did not contain an abstract and/or title. Over the 18-year period, the conference received applications from 128 countries; the majority of applicants were from India (n = 454), United Kingdom (n = 312), Kenya (n = 125), Nigeria (n = 121), or Nepal (n = 100). By region, Asia was the largest source of applicants (n = 992), followed by Africa (n = 961), Europe (n = 598) and Latin America (n = 213) (Table 1).

**Table 1.** Proportion of abstracts across the six regions.

| Region | Nationality | Residence | Fieldwork | % studies based on own fieldwork | % people focused |
| --- | --- | --- | --- | --- | --- |
| Africa | 961 (34%) | 958 (33%) | 1,016 (41%) | 82% | 36% |
| Asia | 992 (33%) | 921 (34%) | 949 (39%) | 83% | 26% |
| Europe | 598 (21%) | 846 (21%) | 216 (9%) | 73% | 27% |
| Latin America | 213 (7%) | 166 (7%) | 222 (9%) | 70% | 32% |
| North America | 86 (3%) | 115 (3%) | 16 (<1%) | 66% | 49% |
| Oceania | 41 (1%) | 55 (1%) | 38 (2%) | 51% | 17% |

Nationality, residence and fieldwork shows the percentage of submissions (after removing those that noted not applicable and not sure) from each region. % fieldwork and % people-focused shows the percentage of submissions, within each region that included fieldwork and a focus on people related values respectively. For the last two columns, submissions were assigned a region based on the nationality of the applicant. Because of different degrees of missing data in individual questions, the sums across columns are not the same.

Noticeably there were very few submissions from North America (n = 86) and Oceania (n = 41). No changes were observed over time in the proportion of applicants from different regions (Fig. S1) and only few, and minor, at the country level (Fig. S2). In terms of country where the study took place, India featured the most (n = 435), followed by South Africa (n = 114), Kenya (n = 110), Nepal (n = 101), and Madagascar (n = 97). Many applicants from Europe and North America worked outside their own region, which was much less the case with students from other regions (Fig. 1). The vast majority of studies were terrestrial (n = 2,393) followed by freshwater (n = 225), marine (n = 177), multiple (n = 119) and coastal (n = 102). Across taxonomic groups, mammals were the most studied (n = 875), followed by plants (n = 470), birds (n = 432), fish (n = 121) and arthropods (n = 89), while potentially-important indicator groups, such as amphibians (n = 58), fungi (n = 10), and lichens (n = 2), were far less represented (Fig. 2).

**Figure 1.**
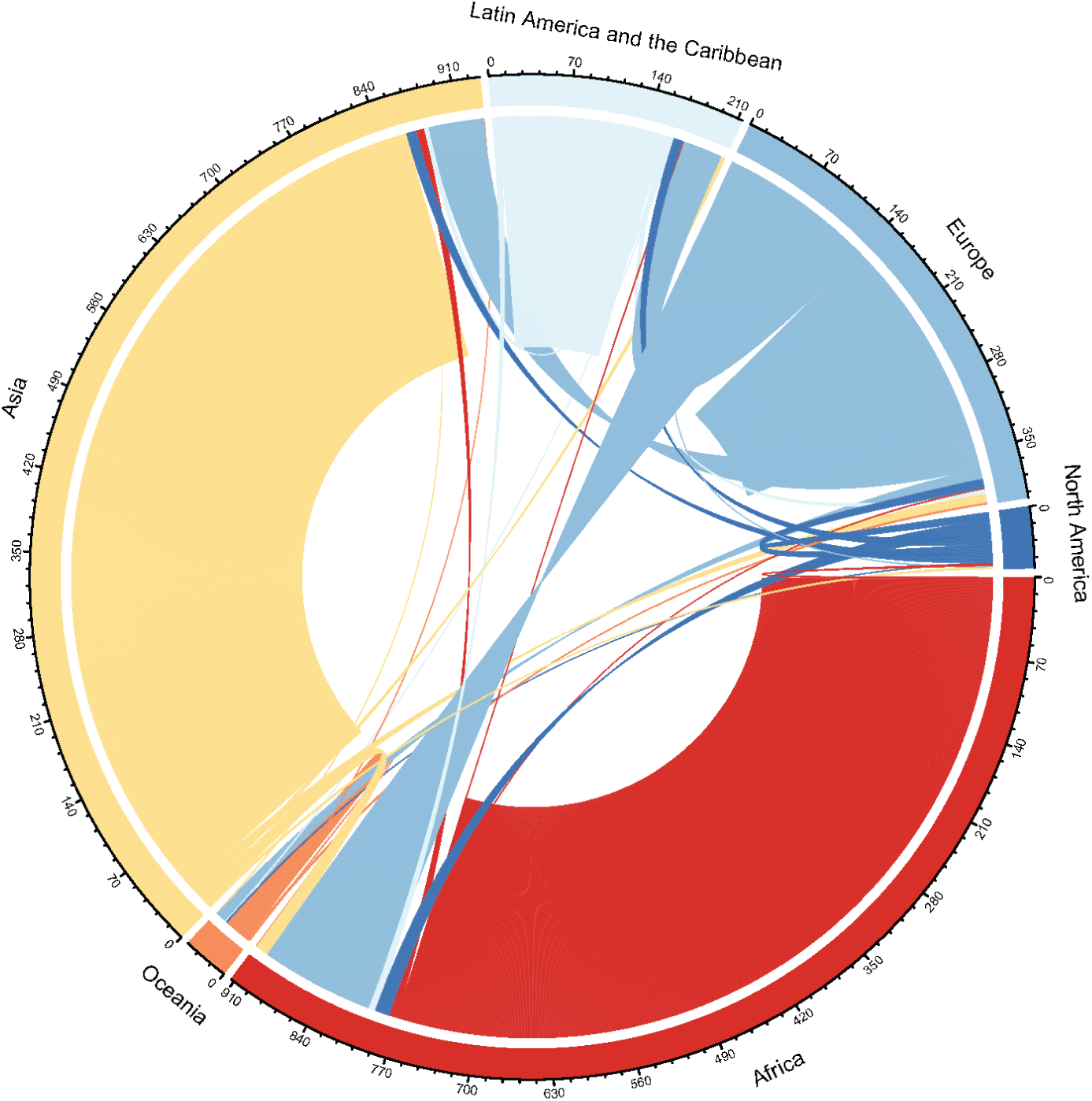
Diagram showing the number of SCCS applicants conducting fieldwork in different regions (the outer ring). The color of the inner (thicker) ring indicates the nationality, grouped by region, of the person conducting the research. The figure shows that there are more Europeans working in Africa, Asia, and Latin America and the Caribbean than there are people from those regions working in Europe.

**Figure 2.**
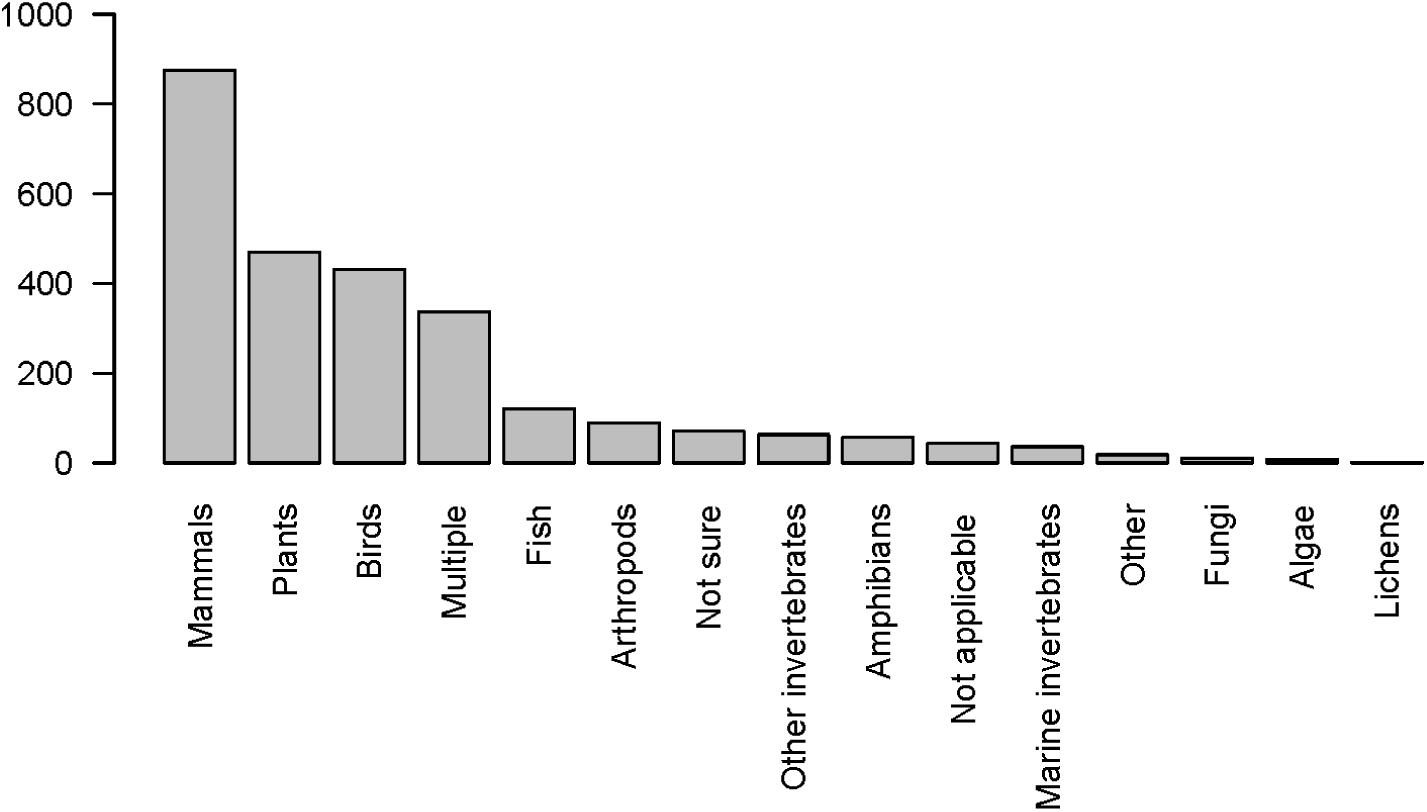
Taxonomic coverage across 2,489 conference submissions (excluding submissions with an ambiguous or no study taxon). Reviewers were explicitly asked to select the main species or higher taxonomic unit of interest. Where other species were described but were not the focus of the study, they have not been recorded.

### 3.2 Framing

On average, 38% (n = 1,003) of studies focused on the state of nature, investigating patterns of biodiversity and processes, followed by 36% (n = 954) addressing pressure to biodiversity, and 26% (n = 671) addressing responses. No changes were observed between 2007 and 2019 in the proportions of state, pressure and response studies (Fig. 3a).

**Figure 3.**
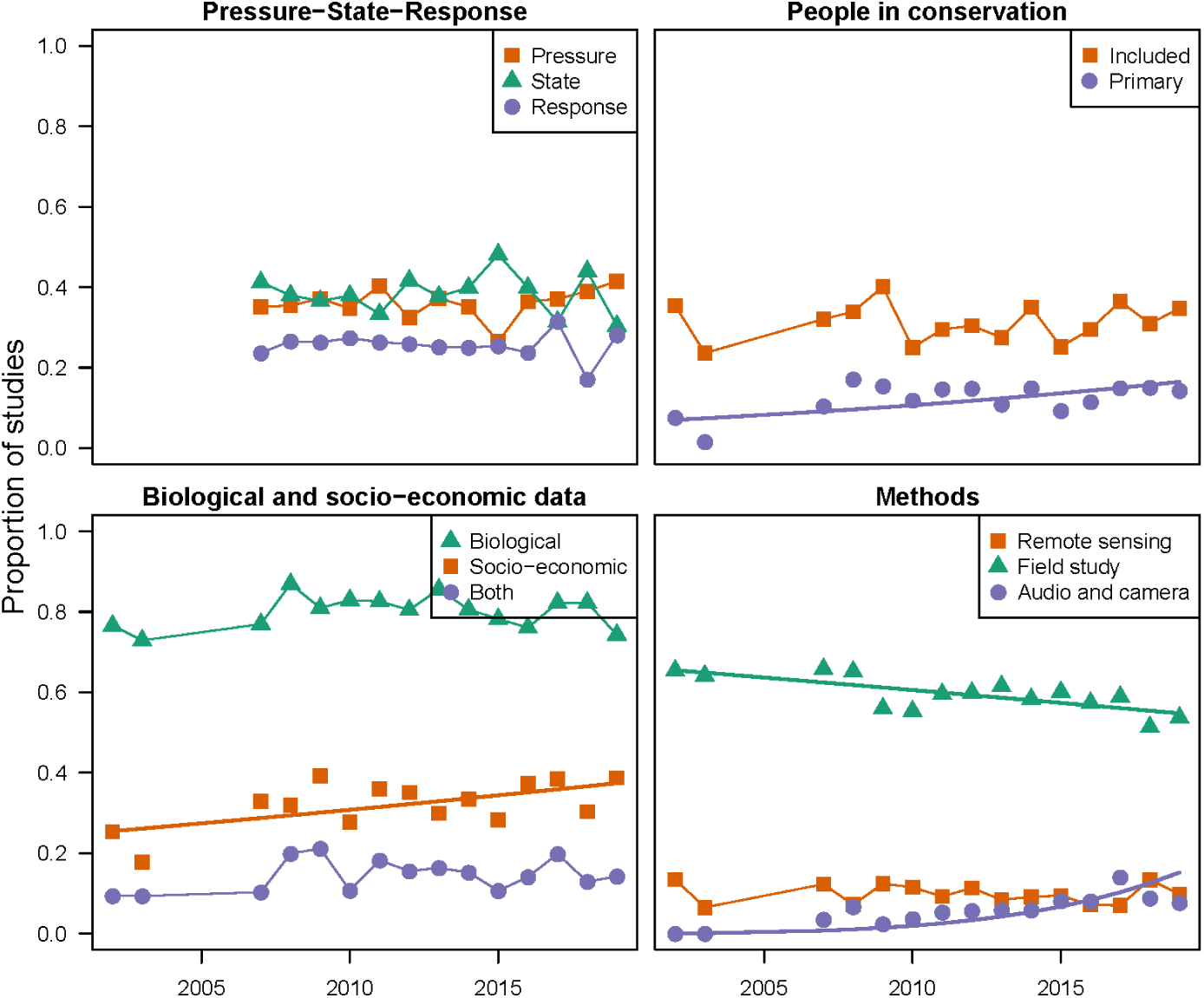
Change over time in the proportions of (a) studies that looked at pressure-state-responses, (b) studies that mentioned the importance of conservation to benefiting people and/or the importance of involving people in conservation decisions, or studies whose primary focus was to understand the values that nature provides to people, (c) studies that including biological data or socio-economic data, or both, and (d) studies using different methods. Studies that noted ‘Not applicable’, ‘Not sure’ or did not provide an answer for the questions involved were not included. Dots are connected where not significant relationship (p ≥ 0.05) was identified while a regression line represents that a significant relationship (p < 0.05) was identified

Of all the submissions, 31.3% (n = 983) mentioned the importance of conservation benefiting people and/or the importance of involving people in conservation decisions, with no change observed over time. While remaining relatively low, in absolute terms (mean = 11.8%) the number of submissions with a primary focus on the value of nature to people increased significantly (z-value = 2.62, p = 0.009, DF = 13) almost doubling from 2002 (estimate = 7.0%) to 2019 (estimate = 16.5%; Fig. 3b).

### 3.3 Data and methods

Most submissions (80%, n = 2,442) contained biological data, while data on socio-economic aspects were less common (33%, n = 998). Only 15% (n = 454) contained both biological and socio-economic data. For biological data and the combination of biological and socio-economic data, the proportions showed no change over time. However, the proportion of submissions including socio-economic data increased over time (estimate = 0.03, S.E = 0.01, p = 0.004, DF = 13; Fig. 3c).

Most of the data, both biological (66%, n = 2,001) and socio-economic 75% (n = 852), were collected by the students themselves. Eighty percent (n = 2,457) of the submissions contained a field-collection element (Table 1) with 74% (n = 2,090) of the submissions covering local-scale studies that looked at one or a few sites, and only 17% (n = 475) of studies investigating patterns at national level, 7% (n = 186) looking at multiple countries, and 2% (n = 66) conducting global analyses.

The methods used to collect biological data remained relatively constant over time and were dominated by field-based approaches, such as transects, plots and trapping (58.4%, n = 1,691). A decrease (from 65.5% in 2002 to 54.8% in 2019) was observed in the use of traditional field-based methods (estimate = -0.026, S.E = 0.006, p < 0.001, DF = 13), and there was an increase (from 0% in 2002 to 15.2% in 2019) in the use of audio and camera recordings (estimate = 0.21, S.E = 0.041, p < 0.001, DF = 13), suggesting a change in approach toward more automated methods, rather than a change away from field-based data-collection (Fig. 3d).

## 4. Discussion

Our results show that the majority of submissions to SCCS were based on primary data collection of biological data from local-scale field studies. These findings suggest a different trend to the concerns raised in previous papers: that there is a decrease in the proportion of field-based studies in the peer-reviewed literature (Carmel et al. 2013; Ríos-Saldaña et al. 2018). Likewise, contrary to the preponderance of researchers from wealthier countries found by reviews of published papers (Amano and Sutherland 2013; Mammides et al. 2016; Martin et al. 2012a), the majority of submissions to SCCS were from Asian and African nationals. These two continents were even more prominent when looking at the countries in which people collected data (Fig 1). For example, citizens from the UK represented the second largest group of applicants, but the UK ranked 15th as a location for fieldwork. The discrepancy, in terms of type and location of studies, between the published literature and submissions to SCCS, highlights a potential barrier in the pathway from fieldwork to publication that warrants further exploration. Furthermore, it suggests that the identified knowledge and data gaps for the tropics, in the published literature (Collen et al. 2008; Mammides et al. 2016; Meijaard et al. 2015), may not only be driven by the lack of research effort and data-collection but also by publication bias. This is of particular concern given many areas in the tropics are of significant biodiversity importance (Brooks et al. 2006; Myers et al. 2000).

There is a need to improve the uptake of studies from the tropics in the peer-reviewed literature to ensure the availability of knowledge and data in conservation research and efforts. This will not only directly benefit the conservation community but will also help ensure a greater diversity in the people and views represented within conservation science. To achieve such improvements, it will be important to support the data-collection-publication pipeline in areas currently underrepresented in the published literature. This may include reduced or waived publication fees (already applied by some journals), as well as language support for non-native English speakers, which is a major barrier in the publication process (Amano et al. 2016). There might also a need for capacity building (Legg and Nagy 2006) and to assist people in scientific writing. For example, in a capacity-building program in Africa run by the Tropical Biological Association, a focus on how to write scientific articles resulted in 87 publications (*Pers obs.* R. Trevelyan). Additionally, while the peer-review process is foundational for the publication of scientific studies, it is not the only way to publish data. An increasing number of preprint services and online data repositories allow for data sharing outside the traditional publication pathway. Similar to the role of GenBank (NCBI Resource Coordinators 2017) in molecular biology, such databases might help to publicize data currently unavailable in the public sphere. However, to be successful, this should be linked to transparent standards (Poisot et al. 2019), formalized method of citing the data-collectors, and must be accompanied by the development of appropriate and fair crediting mechanisms for data collectors by institutions and funding bodies. Additionally, data can represent value, both monetary and cultural, thus where fieldwork is taking place outside the country of the institution the access to data should be accompanied by benefit-sharing (Baker et al. 2019).

Over the 18-year span of the conferences for which we had data, the number of submissions that focused on the value that nature provides to people increased. This corresponds with the emergence within the conservation community of a ‘nature for people’ framing (Mace 2014), which has profoundly influenced the strategies of some of the world’s largest conservation organizations (e.g. Conservation International and The Nature Conservancy; Kareiva et al. 2014). However, this narrative has been criticized as western-dominated (Tallis and Lubchenco 2014) and as describing a polarization not actually found in the conservation community (Sandbrook et al. 2019). In this light, it is interesting that while we observed a significant trend over time, the proportion of SCCS submissions focused on the services and goods that nature provides to people remained low. Thus, our results suggest that while the emphasis on people is a component in conservation, it is by no means dominant. It is possible that our sample, with a majority from lower-income countries, might be less influenced by this trend in conservation than in higher-income regions. North America and Australia which are among the largest contributors to peer-reviewed journal articles in conservation science are for example almost entirely absent in our sample while also among the strongest proponents of a more people focused conservation narrative (Tallis and Lubchenco 2014).

The applications we assessed support suggestions that conservation science is broadening (Teel et al. 2018) by revealing an increase over time in the use socio-economic data. However, the proportion remained relatively low across the 18 years. Additionally, the number of studies integrating both biological and socio-economic data did not increase, with only around 16% of studies combining biological and socio-economic data in the same study. This suggests that while conservation has become increasingly multi-disciplinary, there is still considerable scope for further integration (also see Guerrero et al. 2018). The call for integrating socio-economic perspectives into conservation research is not new (e.g. Adams and McShane 1992), and it is increasingly recognized that both biological and socio-economic perspectives are vital to conservation success (Martin et al. 2016). The continued paucity of socio-economic considerations in conservation science we observed highlights the need to broaden the training of future conservation researchers. This requires university departments and faculties to foster integration and to break down silos between disciplines and departments.

The majority of studies focused on describing biological states or human pressures while only 26% evaluated conservation interventions and solutions. Our results therefore mirror several papers that highlight the lack of studies assessing the impact of conservation responses (Geldmann et al. 2013; Schleicher 2018). While we recognize that an understanding and description of the state of nature and the pressures it faces provides a foundation for developing effective responses, the under-representation of studies assessing the impact of conservation efforts is concerning, given longstanding calls for increasing evidence on the effectiveness of conservation interventions (Pullin and Knight 2001; Sutherland et al. 2004). Assessing the impact of conservation responses is fundamental to improving their effectiveness (Balakrishna 1999; Ferraro 2009) as well as measuring progress towards achieving policy targets (Fisher et al. 2014). It is possible that the complexity of assessing conservation impact (Baylis et al. 2016) is limiting the number of such studies undertaken by students, who are often constrained by time and may lack the experience required to undertake complex impact assessments. However, it is vital that conservation science increasingly addresses this knowledge gap (Baylis et al. 2016; Miteva et al. 2012; Schleicher 2018) to better understand what works, when and why.

By following 18 years of submissions from the oldest student conservation conference, our study provides a unique temporal insight into the work undertaken by successive cohorts and early-career conservation scientists. In including all submissions to give a talk, our sample is not biased by the quality of submissions or by temporal shifts in the preferences of the selection committees but represents the full diversity of students applying for SCCS. Nevertheless, our sample might not represent the wider community as self-selection might exclude some from submitting. As with published studies (Amano et al. 2016; Amano and Sutherland 2013), countries (often former British colonies) where it is more common to communicate in English were disproportionately represented and so the conference doubtless does not fully capture a representative sample of all conservation studies. Moreover, the low proportion of marine studies indicates that SCCS has tended to attract a lower proportion of those working on marine conservation, perhaps due to the organizers having mostly terrestrial experience and networks. Likewise, the dominance of field studies from the tropics in the conference submissions might not reflect a dominance of field studies in general. Rather, it is possible that fieldwork in temperate zones is framed more as ecological research without a conservation focus. Nevertheless, our study suggests that there is an untapped resource in field studies and perhaps more tropical research being undertaken by students from tropical countries than is suggested by the published literature.

## 5. Conclusion

Based on our findings we see an urgent need to make data generated by tropical fieldworkers more widely available, and for increased efforts in examining the impact of conservation interventions. Any approach should ensure the quality of the data as well as equitable access that respects, acknowledges, and includes the data providers. Our results also highlight the need for further integration of disciplines outside biology. Only through combining understanding of both the natural world and human behaviour can we successfully tackle the great challenges facing Earth’s biodiversity, without jeopardizing the sustainable livelihood of our own species.

## Supporting information

SOM Appendix 1 [search_protocol]

SOM Appendix 2 [supplementary results]

## References

Adams, J., McShane, T., 1992. The Myth of wild Africa: conservation without illusions. University of California Press, Berkeley, California, USA.

Amano, T., González-Varo, J.P., Sutherland, W.J., 2016. Languages Are Still a Major Barrier to Global Science. PLOS Biology 14, e2000933.

Amano, T., Sutherland, W.J., 2013. Four barriers to the global understanding of biodiversity conservation: wealth, language, geographical location and security. Proceedings of the Royal Society B: Biological Sciences 280.

Baker, K., Eichhorn, M.P., Griffiths, M., 2019. Decolonizing field ecology. Biotropica 51, 288–292.

Balakrishna, P., 1999. Biodiversity conservation and impact assessment. Current Science 76, 129–131.

Balmford, A., Cowling, R.M., 2006. Fusion or Failure? The Future of Conservation Biology. Conservation Biology 20, 692–695.

Baylis, K., Honey-Rosés, J., Börner, J., Corbera, E., Ezzine-de-Blas, D., Ferraro, P.J., Lapeyre, R., Persson, U.M., Pfaff, A., Wunder, S., 2016. Mainstreaming Impact Evaluation in Nature Conservation. Conservation Letters 9, 58–64.

Bennett, N.J., Roth, R., Klain, S.C., Chan, K., Christie, P., Clark, D.A., Cullman, G., Curran, D., Durbin, T.J., Epstein, G., Greenberg, A., Nelson, M.P., Sandlos, J., Stedman, R., Teel, T.L., Thomas, R., Veríssimo, D., Wyborn, C., 2017. Conservation social science: Understanding and integrating human dimensions to improve conservation. Biological Conservation 205, 93–108.

Brooks, T.M., Mittermeier, R.A., da Fonseca, G.A.B., Gerlach, J., Hoffmann, M., Lamoreux, J.F., Mittermeier, C.G., Pilgrim, J.D., Rodrigues, A.S.L., 2006. Global Biodiversity Conservation Priorities. Science 313, 58–61.

Carmel, Y., Kent, R., Bar-Massada, A., Blank, L., Liberzon, J., Nezer, O., Sapir, G., Federman, R., 2013. Trends in Ecological Research during the Last Three Decades – A Systematic Review. PLoS ONE 8, e59813.

Clark, J.A., May, R.M., 2002. Taxonomic Bias in Conservation Research. Science 297, 191–192.

Collen, B., Ram, M., Zamin, T., McRae, L., 2008. The tropical biodiversity data gap: addressing disparity in global monitoring. Tropical Conservation Science 1, 75–88.

Ferraro, P.J., 2009. Counterfactual thinking and impact evaluation in environmental policy. New Directions for Evaluation 2009, 75–84.

Fisher, B., Balmford, A., Ferraro, P. J., Glew, L., Mascia, M., Naidoo, R., Ricketts, T.H., 2014. Moving Rio Forward and Avoiding 10 More Years with Little Evidence for Effective Conservation Policy. Conservation Biology 28, 880–882.

Geldmann, J., Barnes, M., Coad, L., Craigie, I.D., Hockings, M., Burgess, N.D., 2013. Effectiveness of terrestrial protected areas in maintaining biodiversity and reducing habitat loss, p. 61. Collaboration for Environmental Evidence, Bangor, United Kingdom.

Godet, L., Devictor, V., 2018. What Conservation Does. Trends in Ecology & Evolution 33, 720–730.

Greenwald, N., Dellasala, D.A., Terborgh, J.W., 2013. Nothing New in Kareiva and Marvier. BioScience 63, 241–241.

Guerrero, A.M., Bennett, N.J., Wilson, K.A., Carter, N., Gill, D., Mills, M., Ives, C.D., Selinske, M.J., Larrosa, C., Bekessy, S., Januchowski-Hartley, F.A., Travers, H., Wyborn, C.A., Nuno, A., 2018. Achieving the promise of integration in social-ecological research: a review and prospectus. Ecology and Society 23.

Kareiva, P., 2014. New Conservation: Setting the Record Straight and Finding Common Ground. Conservation Biology 28, 634–636.

Kareiva, P., Groves, C., Marvier, M., 2014. The evolving linkage between conservation science and practice at The Nature Conservancy. Journal of Applied Ecology 51, 1137–1147.

Kareiva, P., Marvier, M., 2012. What is conservation science? BioScience 62, 962–969.

Legg, C.J., Nagy, L., 2006. Why most conservation monitoring is, but need not be, a waste of time. Journal of Environmental Management 78, 194–199.

Mace, G.M., 2014. Whose conservation? Science 345, 1558–1560.

Mammides, C., Goodale, U.M., Corlett, R.T., Chen, J., Bawa, K.S., Hariya, H., Jarrad, F., Primack, R.B., Ewing, H., Xia, X., Goodale, E., 2016. Increasing geographic diversity in the international conservation literature: A stalled process? Biological Conservation 198, 78–83.

Martin, J.-L., Maris, V., Simberloff, D.S., 2016. The need to respect nature and its limits challenges society and conservation science. Proceedings of the National Academy of Sciences 113, 6105–6112.

Martin, L.J., Blossey, B., Ellis, E., 2012a. Mapping where ecologists work: biases in the global distribution of terrestrial ecological observations. Frontiers in Ecology and the Environment 10, 195–201.

Martin, T.G., Burgman, M.A., Fidler, F., Kuhnert, P.M., Low-Choy, S., McBride, M., Mengersen, K., 2012b. Eliciting Expert Knowledge in Conservation Science Obtención de Conocimiento de Expertos en Ciencia de la Conservación. Conservation Biology 26, 29–38.

McHugh, M.L., 2012. Interrater reliability: the kappa statistic. Biochemia medica 22, 276–282.

Meijaard, E., Cardillo, M., Meijaard, E.M., Possingham, H.P., 2015. Geographic bias in citation rates of conservation research. Conservation Biology 29, 920–925.

Meine, C., Soulé, M., Noss, R.F., 2006. “A Mission-Driven Discipline”: the Growth of Conservation Biology. Conservation Biology 20, 631–651.

Miteva, D.A., Pattanayak, S.K., Ferraro, P.J., 2012. Evaluation of biodiversity policy instruments: what works and what doesn’t? Oxford Review of Economic Policy 28, 69–92.

Myers, N., Mittermeier, R.A., Mittermeier, C.G., da Fonseca, G.A.B., Kent, J., 2000. Biodiversity hotspots for conservation priorities. Nature 403, 853–858.

NCBI Resource Coordinators, 2017. Database resources of the National Center for Biotechnology Information. Nucleic Acids Research 46, D8–D13.

Noss, R., Nash, R., Paquet, P., Soule, M., 2013. Humanity’s Domination of Nature is Part of the Problem: A Response to Kareiva and Marvier. BioScience 63, 241–242.

Organisation for Economic Co-operation and Development, 1993. OECD core set of indicators for environmental performance reviews. OECD, Paris, France.

Poisot, T., Bruneau, A., Gonzalez, A., Gravel, D., Peres-Neto, P., 2019. Ecological Data Should Not Be So Hard to Find and Reuse. Trends in Ecology & Evolution 34, 494–496.

Pullin, A.S., Knight, T.M., 2001. Effectiveness in conservation practice: Pointers from medicine and public health. Conservation Biology 15, 50–54.

R Development Core Team, 2019. R: A language and environment for statistical computing. R Foundation for Statistical Computing.

Ríos-Saldaña, C.A., Delibes-Mateos, M., Ferreira, C.C., 2018. Are fieldwork studies being relegated to second place in conservation science? Global Ecology and Conservation 14, e00389.

Sandbrook, C., Fisher, J.A., Holmes, G., Luque-Lora, R., Keane, A., 2019. The global conservation movement is diverse but not divided. Nature Sustainability 2, 316–323.

Schleicher, J., 2018. The environmental and social impacts of protected areas and conservation concessions in South America. Current Opinion in Environmental Sustainability 32, 1–8.

Soule, M., 2013. The “New Conservation”. Conservation Biology 27, 895–897.

Soulé, M.E., 1985. What Is Conservation Biology? BioScience 35, 727–734.

Sutherland, W.J., Pullin, A.S., Dolman, P.M., Knight, T.M., 2004. The need for evidence-based conservation. Trends in Ecology & Evolution 19, 305–308.

Tallis, H., Lubchenco, J., 2014. Working together: a call for inclusive conservation. Nature 515, 27–28.

Teel, T.L., Anderson, C.B., Burgman, M.A., Cinner, J., Clark, D., Estévez, R.A., Jones, J.P.G., McClanahan, T.R., Reed, M.S., Sandbrook, C., St. John, F.A.V., 2018. Publishing social science research in Conservation Biology to move beyond biology. Conservation Biology 32, 6–8.

Tewksbury, J.J., Anderson, J.G.T., Bakker, J.D., Billo, T.J., Dunwiddie, P.W., Groom, M.J., Hampton, S.E., Herman, S.G., Levey, D.J., Machnicki, N.J., del Rio, C.M., Power, M.E., Rowell, K., Salomon, A.K., Stacey, L., Trombulak, S.C., Wheeler, T.A., 2014. Natural History’s Place in Science and Society. BioScience 64, 300–310.

Wilson, E.O., 2017. Biodiversity research requires more boots on the ground. Nature Ecology & Evolution 1, 1590–1591.

