## Supplementary material for "Insights from two decades of the Student Conference on Conservation Science": SOM Appendix 1 [search_protocol]

### SCCS abstract search protocol

All abstracts are placed in folders each corresponding to a year (A-T). the Letters represent a randomized list of the years 2000 and 2019. For most years the folder will contain a series of individual pdfs X_nn, where "X" is the letter of the year and "nn" is the id-number. For some years all abstracts have been compiled into one file. These will have and id number usually in the top right corner of the page.

For both years with individual files and years with one big file the folder contains **all** applications (e.g. people applying to 1) give a talk, 2) present a poster, 3) both talk and poster and 4) just attend). Thus the first task is to look at whether the application has applied to give a talk. **Only abstract with a yes for Talk is to be included.**

| Attendance only | Poster | Talk | Bursary | Intern |
|---|---|---|---|---|
| No | Yes | Yes | Yes | Yes |

Where students have applied to give both a poster and talk – there will be two (usually identical) abstract sections. **Extract the information under the heading TALK**.

**F**or many of the questions there will be drop-down menus with the allowed/appropriate categories. The description of the individual categories (for selected questions) are given in the table below. Should you run in to answers that would require additional categories please e-mail me ASAP to assess whether the table can be amended.

We have abstracts from 2006-2019 – while the format hasn't changed much, a few fields have been modified: Nationality is not always given – where not use "Not applicable". For some years, the abstracts have not been divided in to the four sections: 1) What conservation problem or question does your talk address?, 2) What were the main research methods you used?, 3) What are your most important results? And 4) What is the relevance of your results to conservation? – where that is the case enter the full abstract in the first field: "What conservation problem or question does your talk address?"

"Not applicable" and "Not sure". Most questions will in addition to a set number of categories, have the option to select "Not applicable" (NA) and "Not sure". **Not applicable** is used in situations where the particular question is not relevant (e.g. the "Scale of study" for a laboratory experiment or "Higher order taxonomy" for a study that does not include biodiversity). **Not sure** is used where the text does not allow for extracting the needed information or where conflicting information does not provide an option relating to one of the categories. Binary Yes/No questions will usually also contain options for Not applicable and Not sure.

Excel does not allow for auto-complete as default. However, as you start using the various levels, the system should start suggesting previously used answers.

Where consecutive columns in the excel-spread sheet are linked in themes, the header columns have the same colours. For the questions related to biological data (columns W-AH) and socio-economic data (columns AI-AS) the first columns (W and AI) have a darker shade. If you answer "No" in either of these, just leave the rest of the columns within that field empty (i.e. if there is no biological data you do not need to fill in whether "Biological data primary" or not).

| Theme | Questions | Notes |
|---|---|---|
| Participant | ID | Combines year (coded as a letter so avoid assessors bias based on knowing the year of the abstract) and talk ID. Where each abstract is a separate file the ID is the name of the file (e.g. A_1), where all abstracts are combined in to one file, the ID is the Year-letter "_" followed by the letter, usually given in the top right of the page. |
| Participant | Nationality | Copy and paste field (Not applicable if this is not included in the form or left blank) |
| Participant | Country of residence | This information is not always provided as a separate field, but can usually be extracted from institution (Not applicable if this is not included in the form or cannot be extracted from the form) |
| Participant | Stage in career | Only degree – and not progression (i.e. **PhD** not 3rd year PhD) (Not applicable if this is not included in the form. For students between degrees (e.g. just handed in PhD/master), use the highest completed degree – this obviously only apply where a new degree has not been started. For students now enrolled as degree X but explicitly stating they are presenting work from degree X-n still use the degree at which they are *currently* enrolled.<br>**Levels:**<br>1. PhD<br>2. Master<br>3. Bachelor<br>4. other |
| Research focus | Title | Copy and paste field from **the title given under the Talk section** – not the same as the submission title! |
| Framing | What conservation problem or question does your talk address? | Copy and paste field **(For years where the abstract is not divided in to the four categories – the full abstract is added to this field**) |
| Method | What were the main research methods you used? | Copy and paste field – left empty for abstracts not divided in to four sections. For these the full abstract goes in the field: "What conservation problem or question does your talk address?" |
| Research focus | What are your most important results? | Copy and paste field – left empty for abstracts not divided in to four sections. For these the full abstract goes in the field: "What conservation problem or question does your talk address?" |
| Framing | What is the relevance of your results to conservation? | Copy and paste field – left empty for abstracts not divided in to four sections. For these the full abstract goes in the field: "What conservation problem or question does your talk address?" |
| Research focus | Geographical realm | One or the below categories depending on where the study is located. For theoretical studies, simulations, laboratory etc use "Not applicable"<br>**Levels:**<br>1. Terrestrial<br>2. Marine<br>3. Freshwater<br>4. Coastal<br>5. Multiple<br>6. Not applicable<br>7. Not sure |
| Research focus | Country/countries where study is taking | If the study is taking place in one specific country copy this over. If the study is taking place in Multiple countries use multiple – if the study is of an entire |

| Theme | Questions | Notes |
| --- | --- | --- |
| | place<br><br>[CONTINUED FROM PREVIOUS PAGE] | larger region (e.g. South Asia, Africa, Latin America) the name of the region can also be inputted. If the study is theoretical, laboratory etc. use Not applicable.<br>**Types:**<br>1. Name of country<br>2. Multiple<br>3. Region<br>4. Not applicable<br>5. Not sure |
| Research focus | Scale of study | Studies with a geographical extend are divided in to Local, National, Multi-county (i.e. regional) and Global. The level depends more on the framing than the actual number of sites/cases. If the target is to derive national level inference, even a relatively low number of sites can be considered national.<br>**Levels:**<br>1. Local (1 or a few sites – scope to explore specific process/pattern not to extrapolate to country level)<br>2. National (scope to extrapolate to country level with a sample of multiple sites or general country-level statistics)<br>3. Multi-country<br>4. Global<br>5. Not applicable (i.e. Lab/modelling/theoretical)<br>6. Not sure |
| Method | Laboratory (*in vitro*) | Yes, if any part of study is undertaken in the **lab/controlled environment.** Yes in this box does not preclude that the study also contains a field/modelling element |
| Method | Field (*in situ*) | Yes, if any part of study is undertaken in the **field.** Yes in this box does not preclude that the study also contains a lab/modelling element |
| Method | Remote-sensed/database | Yes, if the study includes any data remotely-sensed or from large-scale data repositories (like national statistics etc.) |
| Method | Modelling (*in silico*) | Yes if any part of study uses **modelling** – modelling is any process involving a computation of data that is not purely to determine statistical certainty of hypothesis or in observed patterns. Yes in this box does not preclude that the study also contains a Lab/field element |
| Research focus | Response-Pressure-State | Assessment of whether the study primarily addresses response (e.g. protected areas, re-introduction), threats/pressures (e.g. climate change hunting etc.), or state of nature/biodiversity (e.g. population structure/abundance/species richness).<br><br>**<u>This variable is hierarchical:</u>** *Response* supersedes *Pressure* which supersedes *State*. – Thus, where a study looks at the effect of protected areas (response) in reducing hunting (a pressure) of lions (state) the study is classified as a “response”, even if it also measures number of poachers and change in lion numbers. Likewise, if a study looks at the threat of invasive species (pressure) on native plants (state) the study is classified as “pressure”. Thus, “state” is used for studies that looks as aspects of biodiversity without relating these to a specific pressure or response.<br>**Levels:**<br>1. Response<br>2. Pressure<br>3. State<br>4. Not applicable<br>5. Not sure |

| Theme | Questions | Notes |
|---|---|---|
| Research focus | Species | If the study is about **<u>one (1)</u>** specific named species this will be recorded (by scientific name if possible, common if not). Where one species is the focus, but other species are mentioned enter the focus species (e.g. the status of Iberian lynx in relation to threats from domestic dogs – in this example lynx should be entered or a network study specifically looking at the role of one species in the network). |
| Research focus | Higher order taxonomy | If species (one or multiple) are recorded, the taxonomic level should also be captured:<br>**Levels:**<br>1. Algae<br>2. Amphibians<br>3. Arthropods<br>4. Birds<br>5. Fish<br>6. Fungi<br>7. Lichens<br>8. Mammals<br>9. marine invertebrate<br>10. other invertebrates<br>11. Plants<br>12. Reptiles<br>13. Other<br>14. Multiple<br>15. Not applicable<br>16. Not sure |
| Framing | Main 5 MEA/IUCN/IPBES threat | If the study addresses one of the five main threat. For other threats use Other – where no threat is mentioned use Not applicable. Where the abstract includes one of the five threats + and additional threat not on the list, use the main threat **not** multiple.<br>**Levels:**<br>1. Land- and sea use change (Habitat loss and degradation),<br>2. Invasive Alien Species,<br>3. Over-exploitation of natural resources<br>4. Pollution and diseases<br>5. Human-induced climate change<br>6. Other<br>7. Multiple<br>8. Not applicable<br>9. Not sure |
| Framing | People-centred conservation / new conservation | Yes, if the abstract mentions/emphasises the importance of conservation to benefiting people and/or the importance of involving people in conservation decisions (e.g. new conservation). This does not imply that the study is primarily framed around the value of nature to people – but that this aspect is articulated (e.g. a study on the diversity of **<u>wild</u>** pollinators articulating that pollination is an Ecosystem service or mapping habitat decline of sacred sites while mentioning the value of these for local culture or a study of deforestation articulating (even if not measuring) the value of forest for resources/water quality) |
| Research focus | Primary objective Benefits to people | Yes, if the primary focus is to understand the value that nature provides to people |

| Theme | Questions | Notes |
|---|---|---|
| Research focus | Biological data | Yes, if study presents any biological data (e.g. species, habitats, genetics or any other data that related to data derived form a biological system) This does not exclude that other types of data (e.g. socio-economic) is included<br><br>NB If you answer No or Not applicable to this leave the remaining biological data columns empty! |
| Research focus | Biological data primary | Yes, if any data included is collected by student/research team and not collated from web-sources or other data-repositories |
| Research focus | Biological data secondary | Yes, if any data included is collated from web-sources or other data-repositories not directly generated as part of the students work or the work of the lab she/he is part of. |
| Research focus | Bioloigical pattern/process studied 1-3 | For reporting the overall pattern/process of biological monitoring/studied<br>**Levels:**<br>1. **Habitat** (Biodiversity is proxied by metrics on the condition of habitat or the environment, rather than directly on organisms themselves. Examples include forest cover, habitat fragmentation, standing biomass, etc. See category 5 for potential exceptions concerning habitat processes.)<br>2. **Individual (pattern)** (Biodiversity data directly measure organisms on the individual level, and concern the condition/state of the target individuals. Examples include the diet of Pallas's cats, the impact of water traffic on the body condition of orcas, the reintroduction of wolves on the stress hormone levels of elk, etc. See category 5 for individual-level cases that should be considered as processes.)<br>3. **Population (pattern)** (Biodiversity data directly measure organisms on the population level, and concern the population abundance and/or structure of target species. Examples include comparing the density of Brazilian tapirs inside versus outside protected areas, the sex ratio of hatchling green turtles over time, the impacts of illegal trade on the population size of Madagascar rosewood, etc. See category 5 for population-level cases that should be considered as processes.)<br>4. **Community (pattern)** (Biodiversity data directly measure organisms on the community level, and concern the structure and/or composition of ecological communities. Examples include the impact of coastal pollution on fish community composition/species richness, the altitudinal gradient of avian functional diversity, the change of plant phylogenetic profile over time, etc. See category 5 for community-level cases that should be considered as processes.)<br>5. **Process** (Biodiversity data concern ecological dynamics that underlie and/or serve as the mechanism for the above-described ecological patterns. Such processes typically involve the interaction of organisms with their biotic and abiotic environment through biochemical, physiological, behavioral, genetic and/or evolutionary pathways, and should typically have manifested/presumed effects on the observed ecological patterns. Examples include the impact of fire disturbance on nutrient cycling processes, the physiological response of mangrove seedlings to increased salinity, the impact of cattle grazing on the habitat use of plains zebras, the influence of climate change on host-parasite relationships between moose and ticks, the demographic impacts of inbreeding in Sumatra tigers due to habitat fragmentation)<br>6. Other<br>7. Not applicable<br>8. Not sure |

| Theme | Questions | Notes |
|---|---|---|
| Research focus | Biological data collection Technique 1-5 | Five columns of methods used to record biodiversity data in the study. Where multiple methods are used these are recorded in separate columns. It is not important whether this was done as part of primary data collection or secondary data collection.<br>**Levels:**<br>1. **FE - Audio-recording** (Field ecology) Any use of audio recording to study behaviour or patterns of one or multiple species. This does not cover where bird-calls are used instead of visual observations.<br>2. **FE - Camera-recording** (Field ecology) any use of camera/video to record behaviour or patterns of species in the wild. This does not include where cameras are mounted on planes/drones.<br>3. **FE - Transect/plot/point count/quadrat** (Field Ecology) Any kind of structured sampling using defined paths or plots.<br>4. **FE - trapping** (Field ecology) Any data collection where individual are trapped (e.g. snares malaise traps)<br>5. **FE citizen science/participatory monitoring** (Field Ecology). Any use of volunteers, citizens or participatory monitoring. The actual task undertaken will often overlap with other categories (e.g. citizens walking transects) in which cases both are entered in separate columns.<br>6. **FE – Other.** Any field based data collection used in the study not captured by the above.<br>7. **G - Environmental DNA** (Genetic). The use of genetic analysis on data collected in the field to identify presence/abundance of species<br>8. **G - Population genetics** (Genetics) The use of genetic material to understand population structures/inbreeding etc.<br>9. **G – Other** (Genetics). Any genetic study not captured by the above.<br>10. **O - Local Ecological Knowledge** (Other). The use of Local Ecological Knowledge to identify features of biodiversity (e.g. people perception of past population sizes, the presence of species)<br>11. **O - Museum Specimens** (Other). The inclusion of data from museum collections – just as with citizen science this will often be associated with another relevant column in which case both are entered in separate columns<br>12. **O - social media** (Other). The use of social media to understand features related to measuring biodiversity – thus people's perception is not included here but covered under socio-economic data.<br>13. **O – Internet/literature** (Other). The use of knowledge from the internet or literature to understand features related to measuring biodiversity – thus people's perception is not included here but covered under socio-economic data.<br>14. **RS - Airplane/LiDAR** (Remote sensing). Any data collected using airplanes or lidar, either human or remotely operated/programmed.<br>15. **RS - Drone** (Remote sensing) any data collected using ground operated flying objects.<br>16. **RS - Satellite** (Remote sensing) any data collected using satellites.<br>17. RS – other (Remote sensing) any remote-sensed data not captured by the above categories<br>18. **Other**. Any biodiversity data collect that does not fall under any of the above categories<br>19. Not applicable<br>20. Not sure |
| Research focus | Biological data technique [Free form] | Record any method or technique described in the abstract free form. Might be useful for text mining or qualitative synthesis. |

| Theme | Questions | Notes |
|---|---|---|
| Research focus | Socio-economic data | Yes, if study presents any data on livelihood issues, economy/finances, attitudes, human behaviours, human behaviour change etc.<br><br>NB If you answer No or Not applicable to this leave the remaining Socio-economic data columns empty! |
| Research focus | Socio-economic data primary | Yes, if any data included is collected by student/research team and not collated from data-repositories |
| Research focus | Socio-economic data secondary | Yes, if any data included is collated from data-repositories not directly generated as part of the students work or the work of the lab she/he is part of. |
| Research focus | Socio-economic objectives 1-3 | Captures the overall objective of the collection of socio-economic data<br>**Levels:**<br>1. Measuring behaviour/activity/direct use (data on what people do, what they collect, where they farm, how they behave/change activity in relation to a conservation intervention etc)<br>2. Measuring values/indirect use of nature/preferences (data on what people feel about nature, environmental values, preferences)<br>3. Measuring attitude to conservation/interventions (data on attitudes to conservation or specific interventions, would include wellbeing assessments if the aim is to evaluate the impact of an intervention on wellbeing)<br>4. Other<br>5. Not applicable<br>6. Not sure |
| Research focus | Socio-economic data collection Technique 1-5 | Method used for collecting socio-economic data.<br>**Levels:**<br>1. Behavior observation (including ethnographic approaches)<br>2. Focus group or participatory approaches (including participatory mapping, participatory video, workshops)<br>3. Log books/diaries<br>4. Questionnaire (a structured survey delivered face to face, online etc)<br>5. Semi-structured interview (often called key informant interview, this is a more flexible approach to data collection than a questionnaire)<br>6. Social media/digital behaviour (e.g. using Instagram posts to infer values, looking at online market places for wildlife species)<br>7. Tracking of humans<br>8. Other<br>9. Not applicable<br>10. Not sure |
| Research focus | Physio-chemical process | Yes, if study presents data on Physio-chemical processes (Climate, weather, topography etc.) |
| Other | Notes | Free form – but should be avoided |
| Other | Flags | The letter(s) of column(s) with issues for potential subsequent check – should be avoided |
