## Supplementary material for "Insights from two decades of the Student Conference on Conservation Science": SOM Appendix 2 [supplementary results]

Supplementary online material appendix 2.

**Notes on years included**

The Student Conference on Conservation Science (SCCS) has run from 2000 to 2019. We have included data from 2002, 2003 and 2007-2019 (14 of the 20 years spanning 18 of the years of the conference). The years 2000, 2001, 2004, and 2005 were excluded because we didn't have all submissions submitted to give an oral presentation for the conference, but only the ones selected for presentation, and thus, could not guarantee no selection bias from the organizers or other sources. For 2006 we had all data, but not the e-mails and therefore never assessed the submissions because of the requirements in the ethics approval. Cooperating

**Table S2.** Kappa scores for questions included in the results section

| Question | Kappa score |
|---|---|
| Country of residence | 0.84 |
| Geographical realm | 0.79 |
| Field work country | 0.78 |
| Scale | 0.70 |
| Laboratory | 0.87 |
| Fieldwork | 0.81 |
| Remote sensed /database | 0.85 |
| Modelling | 0.78 |
| Response/Pressure/State | 0.64 |
| Higher order taxonomy | 0.79 |
| People centred conservation | 0.78 |
| Primary objective: Benefits to people | 0.73 |
| Biological data included | 0.86 |
| Biological primary data included | 0.78 |
| Biological secondary data included | 0.67 |
| Biological data collection Technique | 0.72 |

Moderate covers the interval 0.60 – 0.79, strong covers the interval 0.80 – 0.90.

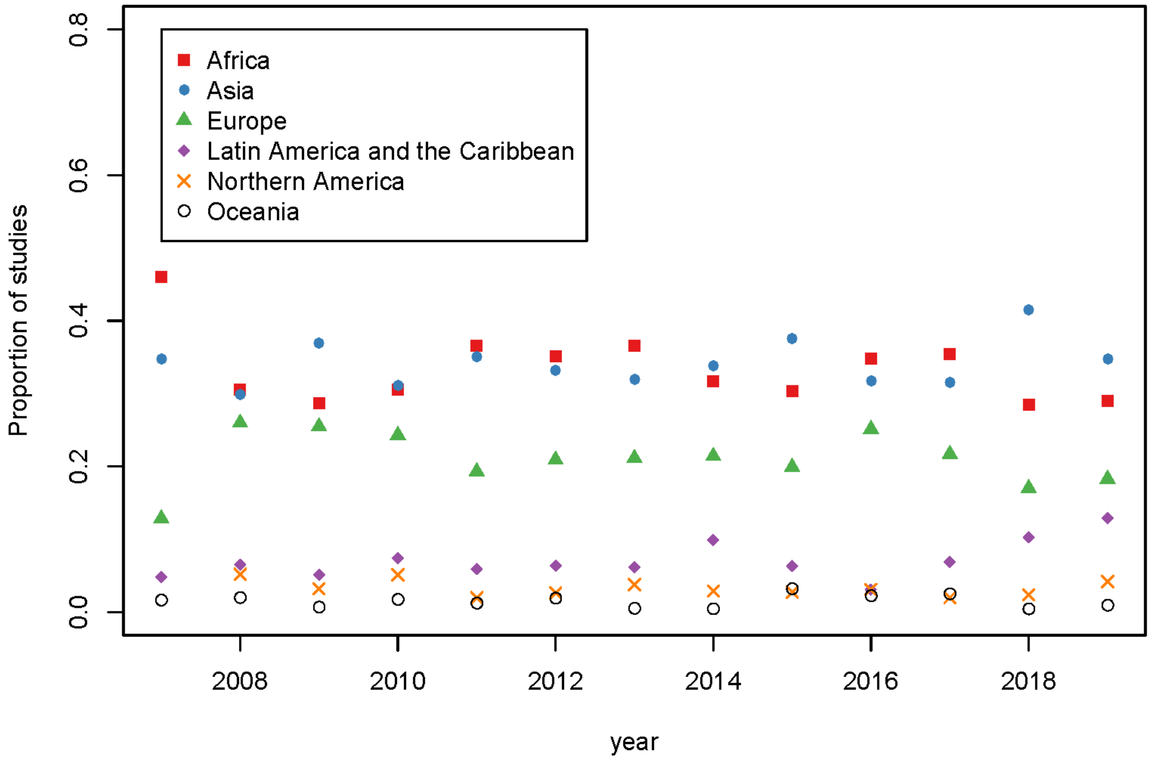


**Figure S1.** Proportion of the six regions over time, showing no trends for any of them.

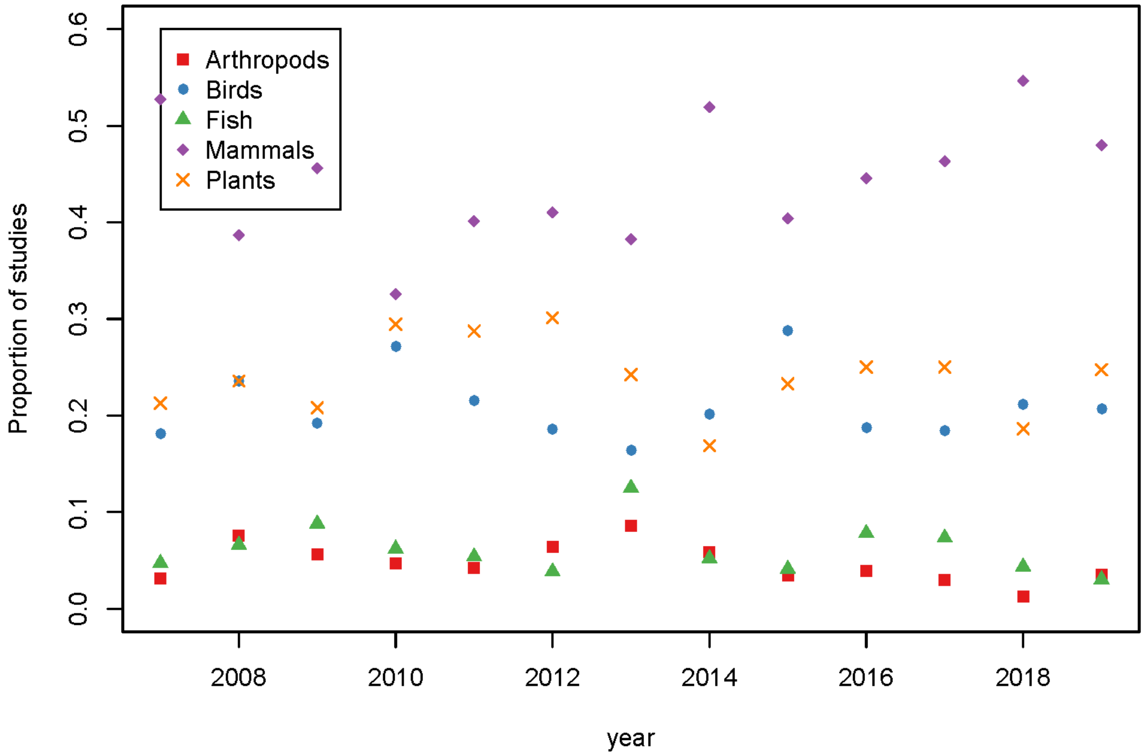


**Figure S2.** Proportion of the top-5 taxonomic groups over time, showing no trends for any of them.

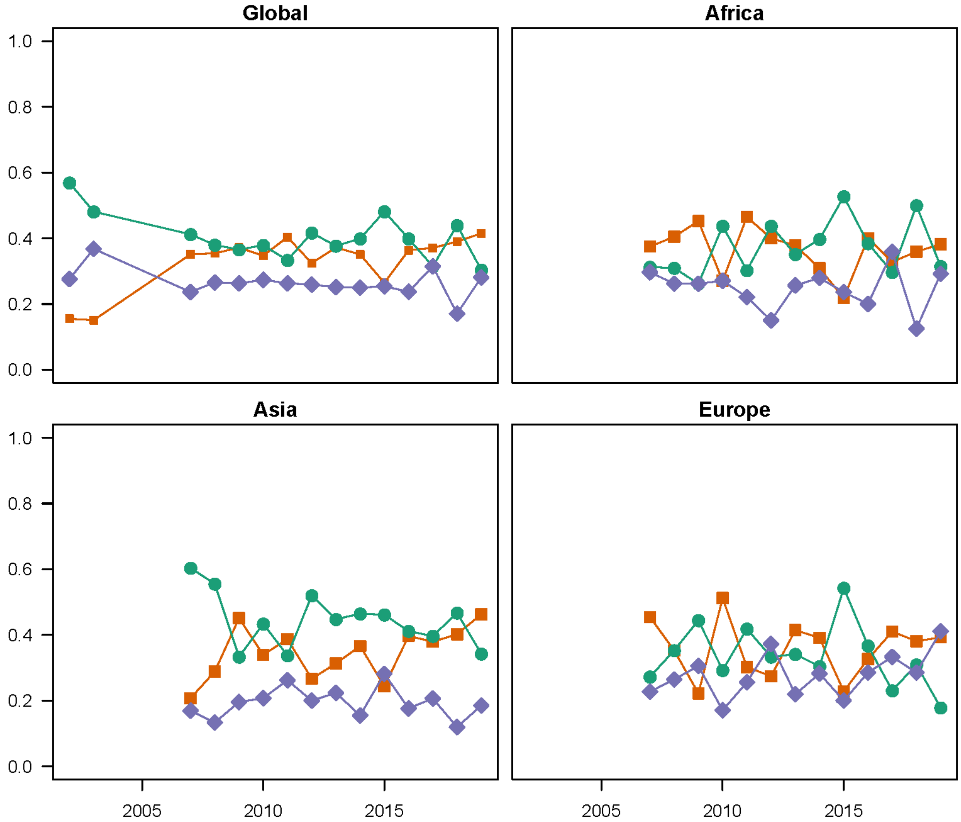


**Figure S3.** Pressure-State-Response for Africa (top-right), Asia (bottom-left), and Europe (bottom-right) which was the three regions for which there was enough data. Green: state, orange: pressure, and purple: response. For none of the regions did we observe any significant changes. For 2002 and 2003, the nationality was not recorded, thus these years are not included in the regional break-downs.

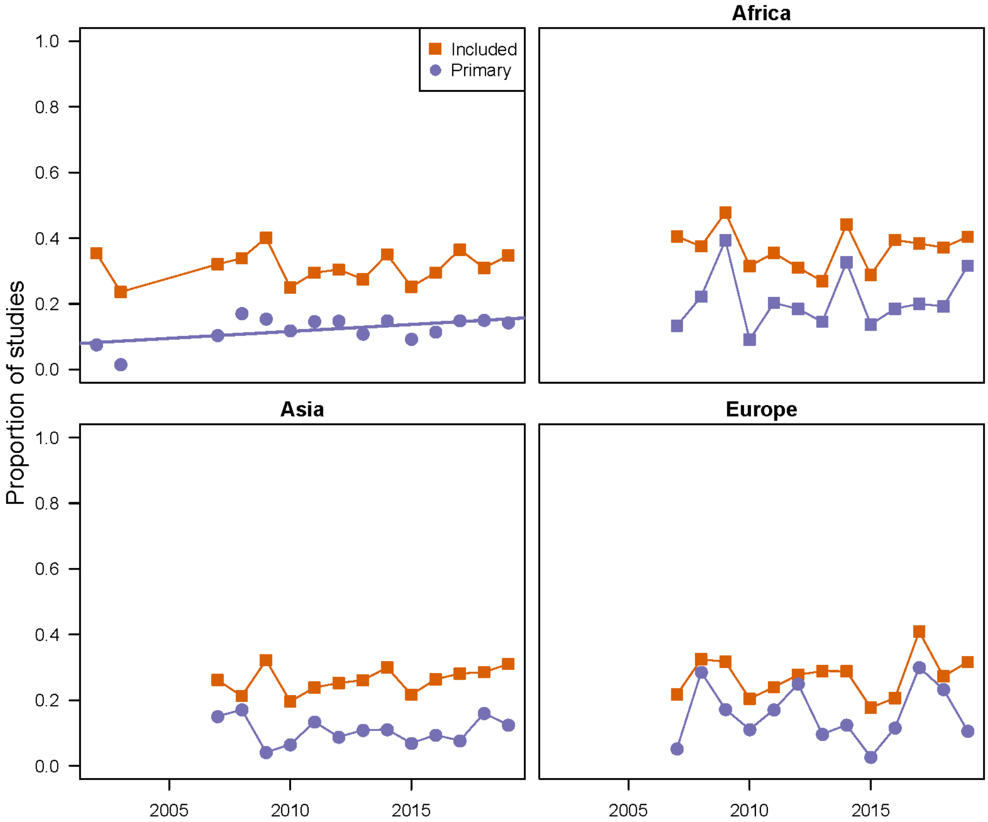


**Figure S4.** People and conservation for Africa (top-right), Asia (bottom-left), and Europe (bottom-right) which was the three regions for which there was enough data. orange: included and purple: primary. For none of the regions did we observe any significant changes. For 2002 and 2003, the nationality was not recorded, thus these years are not included in the regional break-downs.

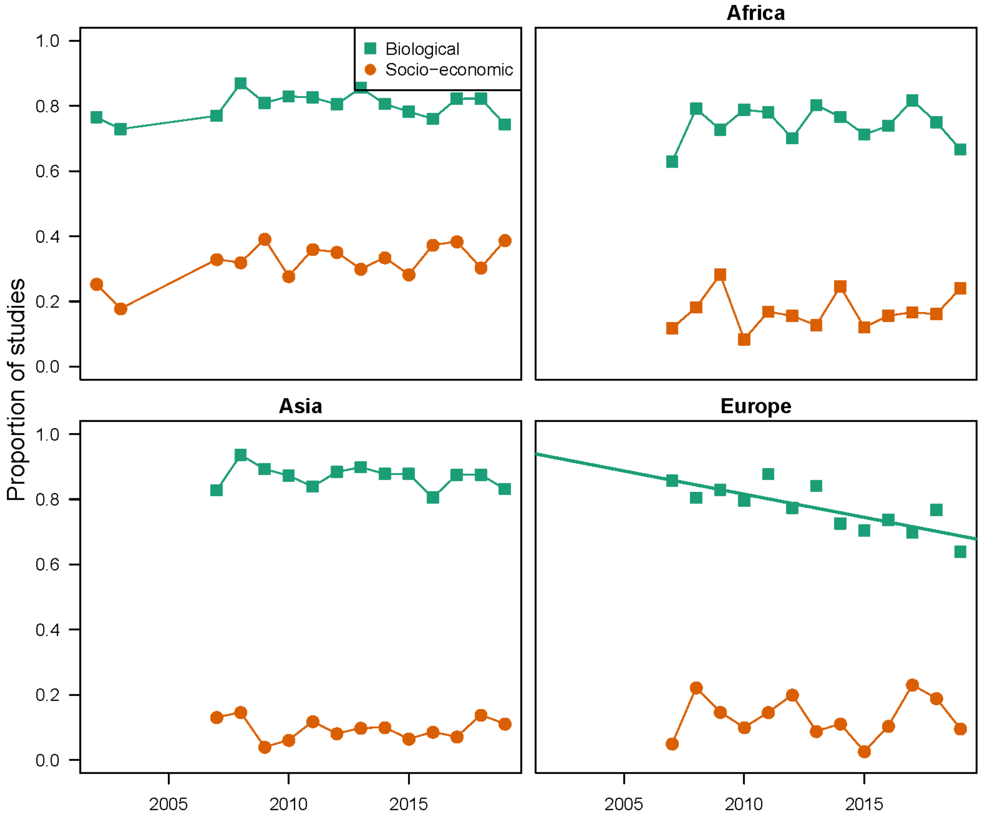


**Figure S5.** Biological and socio-economic data for Africa (top-right), Asia (bottom-left), and Europe (bottom-right) which was the three regions for which there was enough data. Green: biological data and orange: socio-economic data. For Europe, the proportion of biological data decreased over the 13 years for which we had data (). For 2002 and 2003, the nationality was not recorded, thus these years are not included in the regional break-downs.

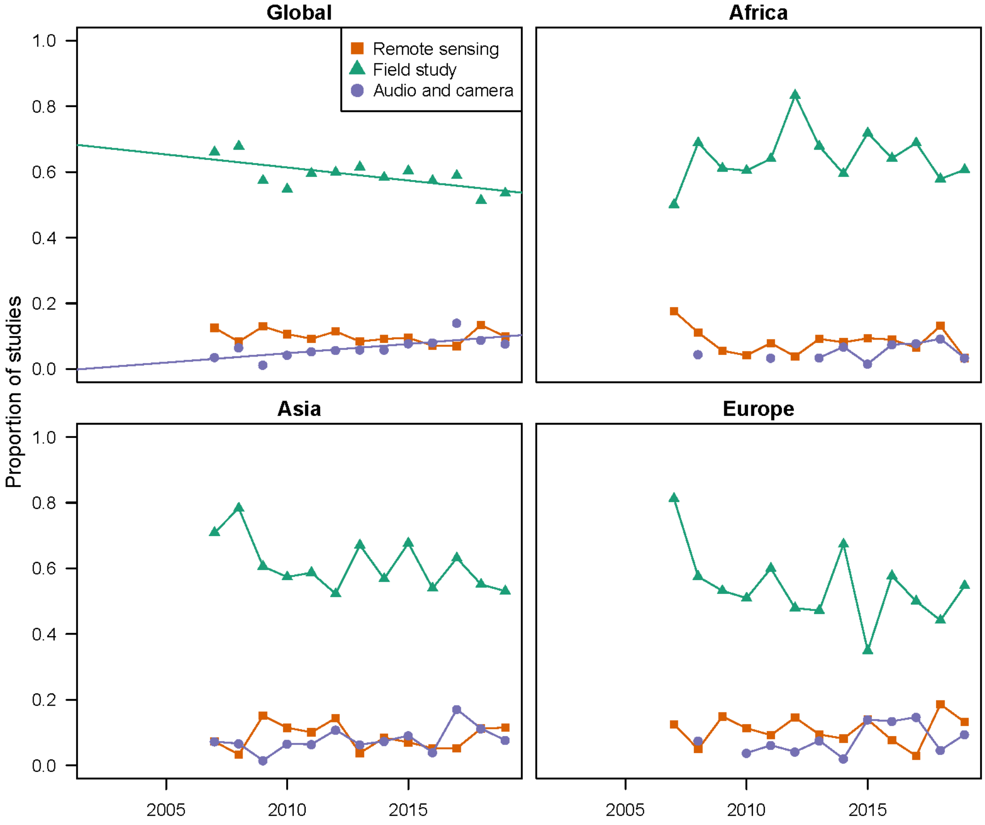


**Figure S6.** Methods per region for Africa (top-right), Asia (bottom-left), and Europe (bottom-right) which was the three regions for which there was enough data. Orange: Remote-sensing, green: field studies, and purple: Audio and camera recordings. For none of the regions did we observe any significant changes. For 2002 and 2003, the nationality was not recorded, thus these years are not included in the regional break-downs.

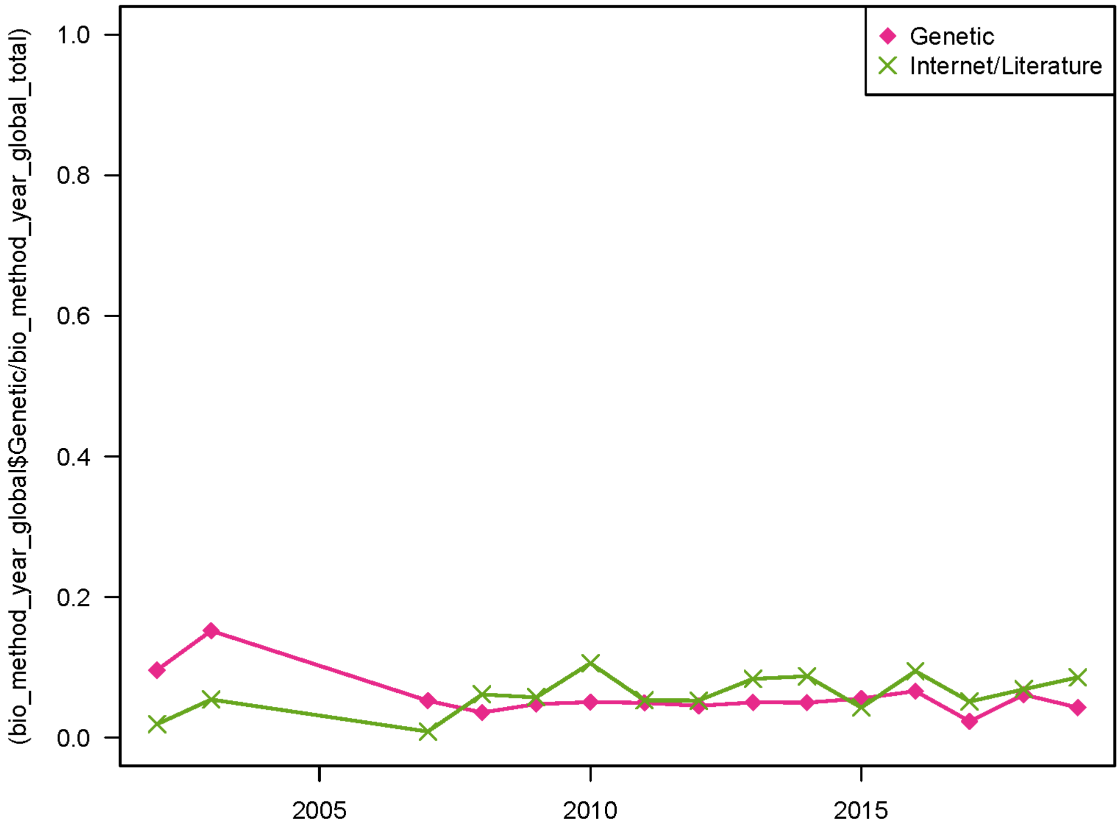


**Figure S7.** The methods not included in the main plot: Genetic (pink), and internet/literature (green).
